# Biological fractionation of lithium isotopes by cellular Na^+^/H^+^ exchangers unravels fundamental transport mechanisms

**DOI:** 10.1101/2022.10.06.510772

**Authors:** Mallorie Poet, Nathalie Vigier, Yann Bouret, Gisèle Jarretou, Saïd Bendahhou, Maryline Montanes, Fanny Thibon, Laurent Counillon

**Affiliations:** Université Côte d’Azur, CNRS, Laboratoire de Physiomédecine Moléculaire (LP2M), Laboratories of Excellence Ion Channel Science and Therapeutics, Nice, France; Oceanography Laboratory of Villefranche (LOV, IMEV), CNRS, Sorbonne University, Villefranche-sur-Mer, France; Université Côte d’Azur, CNRS, Institut de Physique de Nice (INPHYNI), France

**Author notes:** Co-first authors (Mallorie Poet, Nathalie Vigier, Yann Bouret).

## Abstract

Lithium (Li) has a wide range of uses in science, medicine and industry but its isotopy is underexplored, except in nuclear science and in geoscience. ^6^Li and ^7^Li isotopic ratio exhibits the second largest variation on Earth’s surface and constitutes a widely used tool for reconstructing past oceans and climates. As large variations have been measured in mammalian organs, plants or marine species, and as ^6^Li elicits stronger effects than natural Li (~95% ^7^Li) a central issue is the identification and quantification of biological influence of Li isotopes distribution. We show here that membrane ion channels and Na^+^-Li^+^/H^+^ exchangers (NHEs), strongly fractionate Li isotopes. This systematic ^6^Li enrichment is driven by membrane potential for channels, and by intracellular pH for NHEs, where it displays cooperativity, a hallmark of dimeric transport. Evidencing that transport proteins discriminate between isotopes differing by one neutron, opens new avenues for transport mechanisms, Li physiology, and paleoenvironments.

## INTRODUCTION

Evidence for large variations of Li isotopes (reported as δ^7^Li (‰)= ([(^7^Li/^6^Li) / (^7^Li/^6^Li)_lsvec_] – 1) x10^3^), with lsvec as the international standard) in biological samples is compiled in Fig. 1a, as illustrated by the difference between the various organs of a mammal model and its diet (Balter and Vigier, 2014). Li isotopes vary also significantly in modern and fossil carbonate shells, relative to homogeneous seawater (Burton and Vigier, 2011; Penniston-Dorland et al., 2017; Roberts et al., 2018; Vigier et al., 2015), and recent studies (Thibon et al., 2021a; Thibon et al., 2021b) exhibit similar isotopic variations in soft and calcified tissues of marine organisms. Figure 1a highlights that the range displayed by biologic materials is in fact similar as that estimated for the global Earth (~50‰). Another line of evidence of the biological control on Li isotopes comes from studies focusing on the relative distributions and biological effects of pure ^6^Li and ^7^Li related to a therapeutic use (Sherman et al., 1984; Stokes et al., 1982; Ettenberg et al., 2020), suggesting a higher diffusivity of ^6^Li compared to ^7^Li.

**Figure 1:**
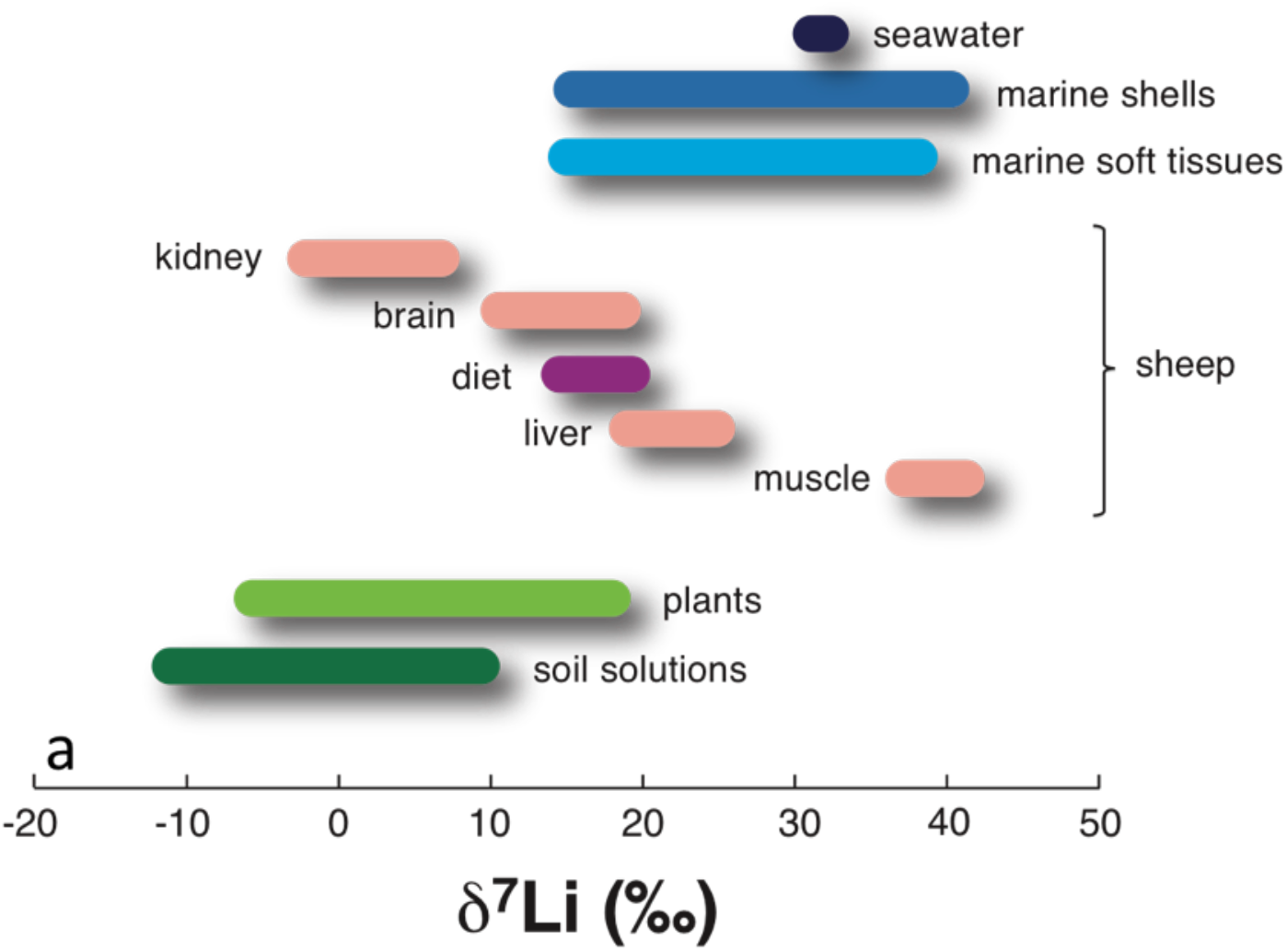
Compilation of published Li isotope compositions measured in biologic materials and their environments. By convention, Li isotope compositions are expressed in δ^7^Li (‰) = (^7^Li /^6^Li) / (^7^Li /^6^Li)_LSVEC_ – 1) x 1000, LSVEC being the international standard). Biologic samples are: (1) in green, terrestrial plants and corresponding soil solutions (Li et al., 2020; Lemarchand et al., 2010) in pink, organs of model mammals (sheep), and corresponding diet in purple (Balter and Vigier, 2014); in blue, living and modern shells produced by marine organisms, and soft tissues of marine species (Pogge von Strandmann et al., 2019; Dellinger et al., 2018; Thibon et al., 2021a), and seawater in dark blue. Note that the ocean is currently homogeneous in terms of Li concentration (26 μM) and δ^7^Li (31.2‰ +/− 0.3‰) (Penniston-Dorland et al., 2017). The range displayed by all the biologic samples is similar as the one estimated for the global Earth (Penniston-Dorland et al., 2017).

The results contained in Figure 1 indicate that Li isotopes accumulate differently in organs and tissues where they can exert distinctive biological effects.

The main way for an alkali cation such as Li^+^ to enter and accumulate in cells is to be transported across their membranes by ion transporters and/or by channels (Hille, 2001; Dubyak, 2004). These proteins are embedded in the membranes of all cells and allow the transport of ions that could not otherwise cross the lipidic bilayer barrier. Ion channels allow the passive flow of ions, following their electrochemical gradients, while transporters use energy-active conformational changes to accumulate ions or other solutes, often in opposition to their concentration gradients. Because cells have a negative membrane potential at rest, Li can enter through various ion channels, but can also be actively translocated through ion transporters. In particular, Li is efficiently transported inside cells by Na^+^/H^+^ Exchangers (NHEs) of the SLC9 gene family, with rates and affinities comparable to Na, their physiological extracellular cation (Pedersen and Counillon, 2019). NHEs are therefore excellent candidates for providing a molecular mechanism for the biogenic fractionation of Li isotopes.

## RESULTS

### Ubiquitous, apical and vesicular NHEs fractionate Li isotopes

We first performed various Li uptake experiments in NHE-deficient fibroblasts (Pouysségur et al., 1984), in which we stably and individually expressed different human NHEs using classical gene transfer techniques and selection (Gao and Huang, 1995; Counillon et al., 1993; Milosavljevic et al., 2014) (SI). The intracellular δ^7^Li value is measured after one minute of Li uptake in cells expressing, respectively, the ubiquitous NHE1, the epithelial NHE2 (Wang et al., 1993) and NHE3 (Orlowski et al., 1992), the vesicular NHE6 (Ouyang et al., 2013) and NHE7 (Milosavljevic et al., 2014) (both expressions directed towards the plasma membrane as in Milosavljevic et al., 2014) as well as in the NHE-null (i.e. PS120, Pouysségur et al., 1984) cells (Fig. 2a and 2b). Compared to the extracellular medium (δ^7^Li=15±0.3‰), all cytosolic contents of NHE-expressing cells show strong ^6^Li enrichments (lower δ^7^Li values), with the highest and the lowest one for NHE3 (by −15.4‰) and NHE7 (by −10‰), respectively (Fig. 2a). The NHE-null PS120 cell line exhibits a minor but non-negligible isotopic fractionation, with an intracellular δ^7^Li value of −4.4‰ lower than the external solution. Thus, NHE-expressing cells, whose exchangers internalize Li in response to cytosolic acidification, display much lower δ^7^Li values compared to equivalent cells, which are not expressing any of those transporters. We therefore conclude that Li isotopes are strongly fractionated by ionic transport through cell membrane. Given that the only difference between these cell lines is the expression of a specific NHE at the membrane, the large isotopic variations exhibited by the NHE-equipped cells are due to active transport performed by the NHEs.

**Figure 2:**
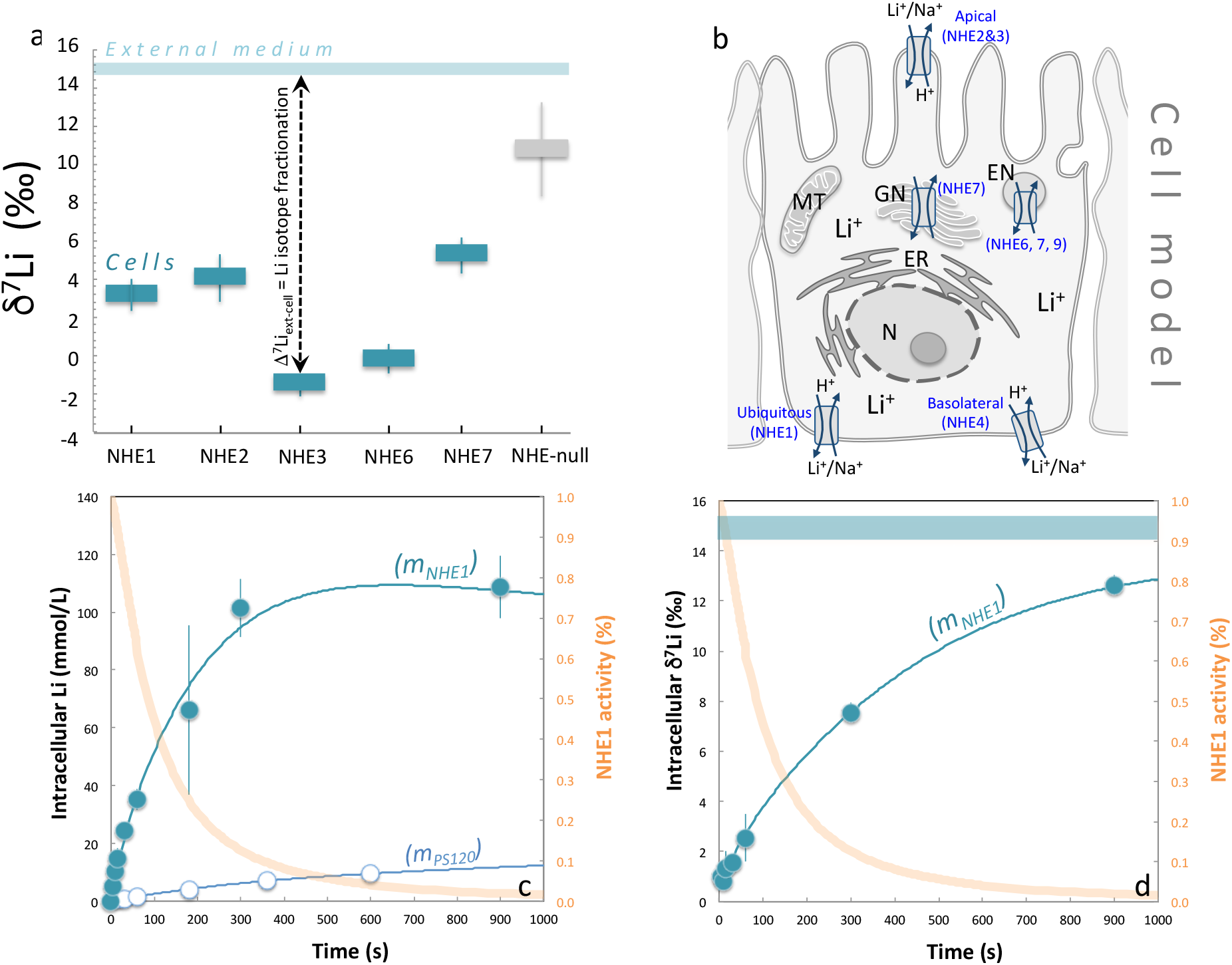
**a. Li isotope fractionation by Na^+^/H^+^ exchangers**. In blue, δ^7^Li values measured in fibroblast cells expressing NHE1, NHE2, NHE3, NHE6 and NHE7, after 1 minute of Li uptake. Li uptake experiments were also performed using NHE deficient PS120 fibroblasts (Pouyssegur et al., 1984) (in grey, NHE-Null). The light blue bar displays the constant δ^7^Li value measured for the external Li uptake solution (15 mM Li). All experiments were performed at 37°C (See Methods in S.I.). Note the large Li isotopic fractionations (Δ^7^Li) between NHE-expressing cells and the external medium solution. **b. Schematic representation of the cellular localization of NHEs**. Li^+^ is an alkali element that is essentially mobile in the outer and inner cellular media. NHE1 is a ubiquitous transporter involved in pH and volume regulation. It is expressed mostly basolaterally in epithelia. NHE2 and NHE3 are apically expressed in epithelial cells, while NHE6 and NHE7 are mostly intracellular and expressed in the Golgi Network and endosomal compartments. N: Nucleus, ER: Endoplasmic Reticulum, GN: Golgi Network, MT: Mitochondria. **c. NHE1 kinetics of Li transport**. In dark blue, Li concentrations measured in NHE1 expressing fibroblast, as a function of the external Li uptake duration. The extracellular solution is the same as in Fig. 1a (15mM Li). In light blue, are reported the same experiments for PS120 cells (NHE-Null). Blue lines (m_NHE1_, m_PS120_) display the transport model results, fitting all data points (see text and SI). In orange is shown the measured NHE1 activity as a function of time (see SI Methods). **d. NHE1 kinetics of Li isotopes transport**. δ^7^Li values (in blue) measured in NHE1 expressing fibroblast as a function of the external Li uptake duration. The blue line (m_NHE1_) displays the NHE1 transport model results, fitting well all data points (see text and SI). The external solution (medium) is the same as in Fig. 1a, with a constant δ^7^Li value over the experiment duration. In orange is shown the measured NHE1 activity as a function of time (see Methods). All experiments are performed at 37°C. At 60 seconds, experiments were reproduced at 20°C (SI).

### Biological Li isotope fractionation is driven by kinetics

We chose NHE1 as a model to quantify biological Li isotopic separation kinetics because it is both well described and present in all eukaryotic cells (Pedersen and Counillon, 2019). For this, we measured fast kinetics of total Li transport in parallel with its isotopic fractionation (SI). In NHE-null cells (PS120), Li enters passively through ion channels, and this can be evidenced and measured using electrophysiological techniques (Fig. S1). Accordingly, Li accumulation followed a two-time scale kinetic (fig. 2c, Table S2), accurately fitted with a Goldman-Hodgkin-Katz equation (Hille, 2001) (describing the reversal potential across a cell membrane), modified to take into account the variations in membrane potential due to the entry of the positive charges provided by Li^+^ itself (SI). This was accompanied, at one minute, with a relatively small isotopic fractionation of −4.4‰ relative to the external medium (Fig. 2a).

In NHE1-expressing cells, rates of Li^+^ accumulation show a short exponential corresponding to pre-steady-state, followed by a quasi-linear uptake corresponding to steady-state behaviour. Finally, Li intracellular concentration stabilizes over long durations (Fig. 2c). Between 5 and 30 seconds intracellular δ^7^Li remained low, at 1.2±0.3‰ on average, compared to 15±0.3‰ for the extracellular medium (Fig. 2d, Table S1). This significant isotopic fractionation (of − 13.8‰), in favour of the lightest ^6^Li isotope, occurred at maximal NHE1 activity and demonstrates the potential of the ubiquitous NHE1 to strongly fractionate Li isotopes, while it maintains intracellular pH. The magnitude and evolution of the cell δ^7^Li value as a function of time rule out a significant contribution of differential sequestration of the two Li isotopes by a hypothetic cytoplasmic component (such as metallothionein in the case of copper isotopes for instance (Albarede et al., 2016; Moynier et al., 2017) and are consistent with the fact that monovalent cations such as Li^+^, Na^+^ and K^+^ are extremely mobile within cells (Hille, 2001).

### pH induced and dose-response of Li isotope fractionations reveal cooperativity

As NHEs interact with both the external and cytosolic medium to transport H^+^ and ions, the corresponding transport rates depend on the external solution concentrations of these ions, as well as on the intracellular pH. Therefore, measuring those rates at different ionic concentrations and pH (i.e. dose-responses) provides valuable information, which are complementary to those provided by kinetics.

We subsequently determined the Li isotope fractionation between the external medium and cells (Δ^7^Li) - after 1 minute - at different H^+^ intracellular concentrations (different cytosolic pH). As NHE1 kinetic is allosterically regulated by cytosolic pH (Lacroix et al., 2004), we expected the Δ^7^Li value to be larger in magnitude at more acidic pH, when the transport activity is stronger (SI). In contrast, at higher intracellular pH, Li isotope fractionation was expected to be smaller, because NHE1 is slower, and passive transport by ion channels would take over. This is indeed what we observed (Fig. 3a&b, Table S5): the largest differences between cells and external solution (Δ^7^Li =−12.5±0.9‰) are measured for the lowest intracellular pH (Fig. 3b). As intracellular pH dictates NHE1 rates, these results indicate that Li isotopic fractionation is kinetically driven by intracellular proton.

**Figure 3:**
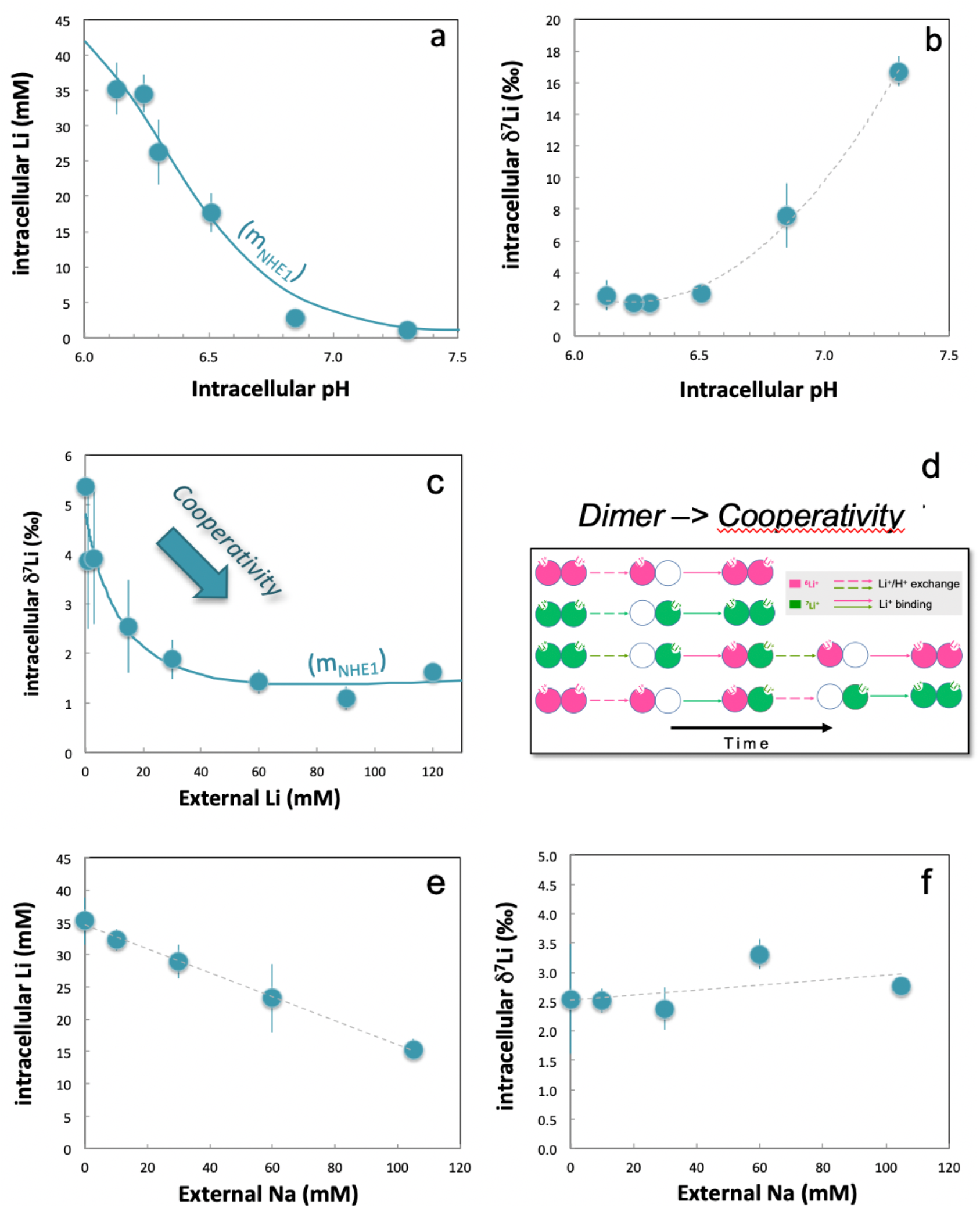
Dose responses for Li isotopic fractionation by NHE1: **a&b. Intracellular pH dependency on Li isotopic transport by NHE1**. Cell Li concentration (a) and δ^7^Li value (b) as a function of intracellular pH. Li uptake was performed during 1 min, at various cytosolic pH for NHE1-expressing fibroblasts. See Methods in SI for details on cell pH control and calibration. The Li transport and associated Li isotope fractionations are maximum at the lowest intracellular pH when NHE1 is at its maximal rate. Total Li uptake was fitted using the MWC equation for a dimeric NHE1^16^, with K_h_ = 0.17 10^−7^M, K_l_ = 36 10^−7^ M, L_0_ = 878.6 ± 35.3 (4.02%), Vmax = 55.4 ± 0.83 (1.51%) and R^2^ = 0.981. **c&d. Dose response for extracellular lithium and cooperative behaviour**. δ^7^Li values of NHE1-expressing cells were measured after one-minute incubation at different extracellular Li concentrations (SI). Note the steady decrease of δ^7^Li as extracellular Li increases, highlighting the unexpected cooperativity of the Li transport process. d. Simplified scheme illustrating how a dimeric NHE1, binding external two ^6^Li^+^ (in pink), two ^7^Li+ (in green) or a mixing of both, before exchanging them with H^+^ will favour the lighter isotope transport. **e&f. Impact of external Na concentration on Li isotopic transport by NHE1**. Intracellular Li concentration (c) and δ^7^Li value (d) as a function of external Na concentrations. The presence of Na in the external medium decreases the Li transport, due to competition effects. In contrast, the cell δ^7^Li value evolves little, showing a slight increase only.

Experiments performed at various extracellular Li concentrations, from 1 to 120 mM, yielded an unexpected result (Table S3). Total Li uptake displayed a typical Michaelis-Menten saturation, with a fitted Km value of 9.94±0.71 mM (r^2^=0.99) in accordance with published data (Table S3). In contrast, the isotopic ratio of transported Li exhibited interesting variations, starting from a δ^7^Li value of 5.4‰ (for an extracellular Li concentration of 0.3 mM), and then steadily decreasing to reach a value around 1.4‰, for media Li contents greater than 60 mM (Fig. 3c). This decrease reveals that, as the external Li concentration increases in total, ^6^Li enhances its own transport against ^7^Li. Such a result is the hallmark of positive cooperativity, which occurs here for the transport of an isotope. As NHE1 is a dimer that displays cooperativity for internal H^+^(Lacroix et al., 2004), cooperativity for external cations at steady-state was expectable, but has been so far impossible to detect, with the only previous evidence coming from daring pre-steady-state experiments (Otsu et al., 1989). Here, this phenomenon is unravelled by the unprecedented high resolution given by Li isotopic measurements by MC-ICP-MS (SI).

### Mathematical derivation of Li isotopic fractionation by a dimeric exchanger

In order to reach a fine mechanistic understanding of our experimental results we developed a mathematical framework for isotopic transport, based on the kinetics of Li and Na transport by NHE and ion channels. ^6^Li and ^7^Li differential transport in NHE1-expressing cells can be calculated by summing the rates of their passive entry through channels, with those of NHE active electroneutral transport. The negative membrane potential triggers the electro-osmotic accumulation of Li species. This is then counterbalanced by the main electrogenic ions, namely by an exit of K^+^ and an entry of Cl^−^ proportional to their respective permeabilities, which, in turn, modify the membrane potential according to the net charge balance. The corresponding Goldman-Hodgkin-Katz flux equations derived from (Hille, 2001) are developed in the SI.

As NHE1 is electroneutral, its transport rate only depends on the inner and the outer transported cations. On each protomer, we formalized NHE1 transport in three simplified steps, in accordance with the Na^+^/H^+^ exchange elevator model for ion translocation (Coincon et al., 2016; Dong et al., 2021) : (1) Li^+^ and H^+^ ions bind NHE in fast pre-equilibria (2) a translocation mechanism occurs that exchanges the NHE-bound ions, (3) NHE is recycled to the initial state, concomitantly extruding the proton to the outer medium and Li^+^ to the inner medium. We described each elementary transformation by a microscopic kinetic constant of the appropriate order and minimally simplified the overall scheme, assuming that the transformations occurring within the transmembrane part of the transporter form fast pre-equilibria.

The kinetic isotopic effect on each microscopic constant produced different rates for the two isotopes. We next introduced the dimeric NHE using the “flip flop” mechanism (Lazdunski et al., 1971) described by Otsu et al. (Otsu et al., 1989) in which each protomer alternatively performs one elementary exchange. Thus, a protonated NHE1 sequentially binds a ^6^Li or a ^7^Li on a first protomer, and then a second ^6^Li or ^7^Li on the other protomer before it can start to exchange ions (Otsu et al., 1989) (Fig. 3d). Following this, a dimer having bound two ^6^Li (E_66_) has the shortest and fastest pathway to regenerate E_66_ after having released one ^6^Li (Fig 3d and SI). In contrast any conversion of a dimer having bound two ^7^Li (E_77_) into E_66_ requires a larger number of steps. Without introducing any additional *ad hoc* change (e.g. affinity or conformation), this combinatorial effect leads to a first come-first serve mechanism, in which ^6^Li transport enhances itself, which is what we observe (Fig. 3c).

A complete derivation for isotopic transport is given in the SI. As in previous modelling studies of ionic transport (Bouret et al., 2014), this approach is mechanistic as it is data driven, and based on physical laws and kinetics of ion transport. It is of note that the resulting set of equations fits (i) the total Li intake (Fig. 2c), (ii) the isotope fractionation kinetics (Fig. 2d) and (iii) the cooperativity of ^6^Li transport within the dimer (Fig. 3c&d), which supports the relevance of our approach. Taken together, modelling NHE Li isotopic transport shows both how the H^+^ gradient drives the isotopic separation and that the sequential binding steps of a dimeric NHE1 trigger positive cooperativity when the external Li concentration increases. As expected, it also shows that the isotope fractionation reaches a saturation value at high external Li concentrations (>>100mM). The model determines, for NHE1, a maximal isotopic fractionation between cell and solution of −21.3‰ (±-2‰). This calculated value is slightly greater than what we found after a few seconds of activity, pointing to the very rapid turnover of NHE1, which changed the cell pH after 5 seconds only. Taken together, NHE performs a fractionation that is 6 times more efficient than electro-osmotic fractionation by ion channels (2-3‰) at the same temperature (SI).

We also show that NHE1 mediated-isotope fractionation is very moderately decreased when Na^+^ concentration increases (Fig. 3e & f, Table S4). This is an interesting difference compared to the observed effect of the inhibitor Cariporide (Scholz et al., 1995), which limits the total Li NHE1 transport by competition, but cannot affect the measured Li isotopic fractionation occurring when Li is bound and transported (SI). Na^+^ behaves differently as it is able both to bind and be translocated, such as Li^+^. Hence in a solution containing both extracellular Li and Na, the dimeric NHE1 is able to bind two Na^+^ ions (in this case no Li^+^ transport occurs), two Li^+^ ions (in this case NHE1 fractionates as described above), or one Na^+^ and one Li^+^ (and in this case NHE behave as a monomer with respect to Li^+^). Consequently, when Na^+^ concentration increases, the proportion of “mixed” versus “fully bound to Li” NHE increases as well, thereby decreasing the cooperative effect and slightly the intracellular δ^7^Li value (Fig. 3f).

## DISCUSSION

Overall, our study demonstrates that ion transport proteins are able to discriminate atoms by their difference in the number of neutrons. In order to translocate between local potential grooves within the protein structure, which has recently been solved for NHE (Dong et al., 2021), Li^+^ ions have to cross an energy barrier. The lighter ^6^Li isotope - which has a greater vibrational velocity, and therefore energy, has a smaller barrier to cross (Fig. 4). This yields greater transport rates that account for the isotopic effects observed in this study. Another important implication for transport is that the Li isotopic analyses of dose responses reveal the positive cooperativity for ^6^Li^+^. Hence, isotopic transport can be established as a novel experimental method to probe the molecular mechanisms of ion channels and transporters with an unprecedented resolution.

**Figure 4:**
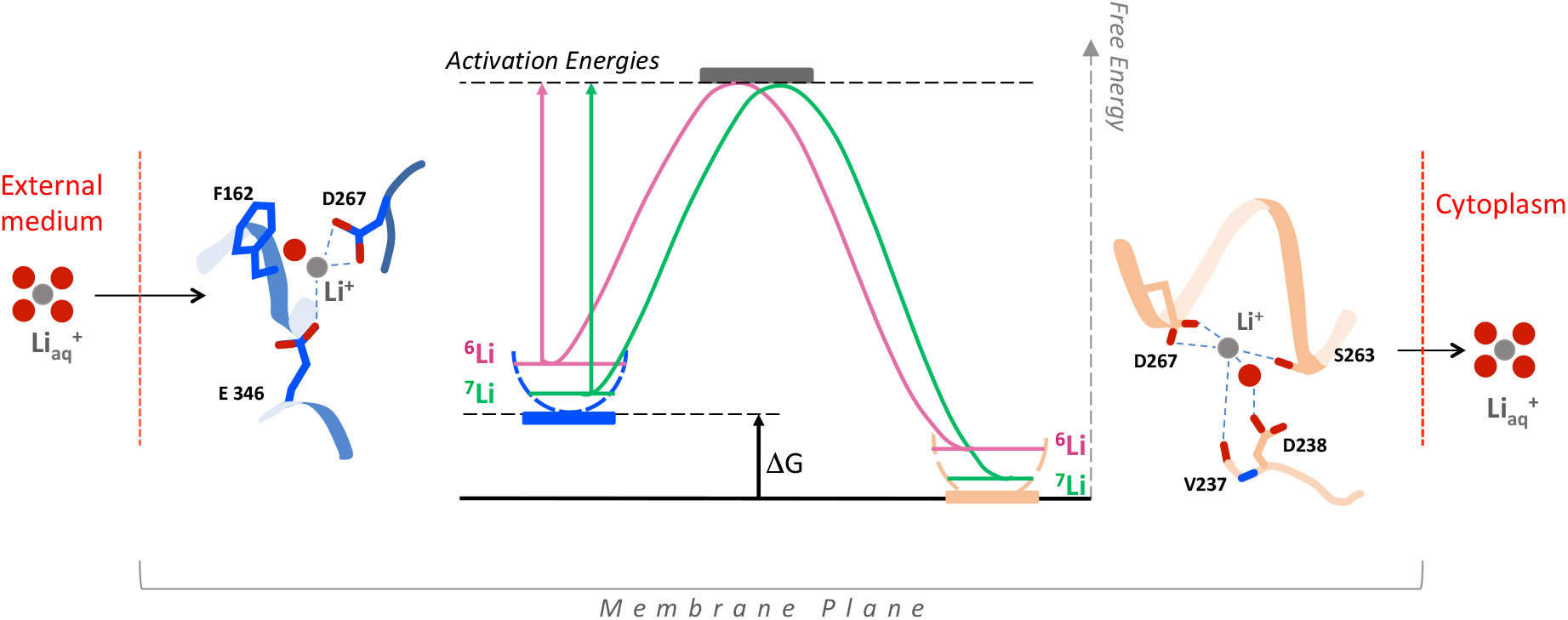
Schematic illustration of the NHE1 isotopic effect. During ion translocation between outward and inward-facing local energy grooves, Li^+^ ions have to cross an activation energy barrier that is smaller for the lighter ^6^Li^+^ (pink) than for the slower and heavier ^7^Li^+^ ion (green). The schemes for Li coordination are drawn from the recently published NHE1 cryo EM structure (Dong et al., 2021). The outward facing conformation (blue) is extrapolated from the NHE1 guanidinium binding site, the cytosol facing conformation (light brown) corresponds to the thallium binding, as described both in (Dong et al., 2021).

The kinetic results presented here show that Li isotopic fractionation persists as long as ion transport is operating. This has interesting potentialities for organ physiology. Indeed, in kidneys, apical NHEs (NHE2, NHE3) are constantly active to reabsorb Na^+^ and extrude H^+^ in order to catalyze bicarbonate reabsorption. Thus, our data may very well explain the range found experimentally in sheep kidney (Fig. 1a). As neurons, astrocytes and the blood brain barrier undergo an intensive ion transport activity, our results also provide clues for the enhanced behavioural effects of ^6^Li that is reported in the literature in model animals. In conclusion, measuring Li isotopes and their transport in soft tissues could be a new field of interest to explore kidney, brain - and possibly other organs function or pathological evolution (Prasad and Rao, 2018).

Implications in geoscience can be developed as well. Indeed, ^6^Li^+^ enrichments (low δ^7^Li values) have been observed in marine biogenic carbonates, especially over the last 70 Ma period, and during short-term climatic or mass extinction events (Hathorne and James, 2006; Kalderon-Asael et al., 2021; Misra and Froelich, 2012; Pogge von Strandmann et al., 2019; Pogge von Strandmann et al., 2020; Washington et al., 2020). In vivo, CaCO_3_ mineral precipitation in cells produces large amounts of H^+^ ions that are eliminated in part by NHEs, which in turn enrich the precipitate in ^6^Li^+^. Ion transport activities are influenced by the external conditions (pH, temperature, Li concentrations). Consequently, our results provide an alternative interpretation of fossil data, more closely related to biological activity variations. Hence the relationship between climate and continental chemical weathering (which removes CO_2_ from the atmosphere and delivers aqueous Li to the ocean) was likely much less strengthened than initially proposed (Caves Rugenstein et al., 2019). Finally, as the genes encoding ubiquitous Na^+^/H^+^ exchangers are thought to have encoded one of the primary ion transport structures of the first protocellular systems (Lane and Martin, 2012), our study also suggests that biological Li isotopic fractionation could be a hallmark of life.

## Supporting information

supplementary tables, figure and full mathematical model

## LIMITATIONS OF THE STUDY

While we clearly found that the background low isotopic fractionation follows a mechanism mediated by ion channels, we did not molecularly identify these channels one by one in the used cell line. We considered that at this step it was too much out of the scope of the present work which mainly focused at exploring how ubiquitous NHE exchangers were responsible for lithium biological fractionation.

## ACKNOWLEDGMENT

The authors acknowledge Philippe Telouk for support with MC-ICP-MS at the CNRS-INSU Facilities located at ENS-Lyon (France). Philippe Campadonico and Jean-Jacques Pangrazi and Nina Milosavljevic (University of Manchester) for critical reading of the manuscript, Abby Cuttriss (University Côte d’Azur Office of International Scientific Visibility) for correcting and improving the English version of the manuscript, to Fadia Boudghene-Stambouli (IRIC Montreal) for her help as a master student and to Michel Tauc (LP2M) for fruitful discussion.

## AUTHOR CONTRIBUTIONS

LC and NV led the project. MP and LC performed experiments with cells. NV, FT, MM, and MP performed sample preparation and Li purification. NV performed Li isotopic ratios and Li concentration measurements. MP performed Li measurements by AAS. YB modeled the dataset. LC and NV wrote the original draft of the manuscript. All authors contributed to review and editing the manuscript. All authors have given approval to the final version of the manuscript

## METHODS

### Methods used for quantifying Li Isotopes transport by NHEs

All experiments were conducted on cells acidified using the ammonium prepulse technique (Boron et al., 1976) in the absence of extracellular sodium, so that the NHEs could only use Li^+^ as a coupling cation to extrude intracellular H^+^ ions (Milosavljevic et al., 2015). Uptake was then stopped by 4 rapid rinses at 4°C, in order to eliminate residual extracellular Li. Cells were lysed in concentrated nitric acid. After Li separation in a clean laboratory using cation exchange resin columns, Li isotopes were measured using the MC-ICP-MS Neptune *Plus* (Thermofisher). More details on these techniques are reported below

### Cell culture

Fibroblasts from the PS120 cell line (NHE-null cells Pouyssegur et al., 1984) were used either as control cells or stably transfected with the pECE expression vector containing NHE1, NHE2 or NHE3 cDNAs (Counillon et al., 1993) to measure the isotopic fractionation mediated by these exchangers. PS120 cells stably transfected with NHE6 (pIRES neo vector, Clontech) or NHE7 (Milosavljevic et al., 2014) (pIRES hygro vector, Clontech) were used to express these NHEs at the plasma membrane by repeated acute acidifications in the presence of 120 mM extracellular NaCl, as described in (Milosavljevic et al., 2014). All cells were cultured at 37°C in a humidified 5% CO_2_-95% air atmosphere.

### Measurement of Li uptake

All experiments were performed without CO_2_ and bicarbonate to ensure that the only pH regulating transporters are NHEs. Cells seeded on 24-well plates were acidified using the NH_4_^+^ loading technique[Boron76]. Measurements were performed at 37°C by incubation in the uptake medium with various concentrations of LiCl (Sigma Aldrich) and the impermeant choline-chloride cation to maintain osmolarity. After the indicated durations, uptake was stopped by four rapid rinses in ice-cold PBS. Cells were solubilized in nitric acid (trace metal grade, Fisher Scientific) and aliquoted for Li concentrations and for Li isotope analyses. Total intracellular Li^+^ concentrations were measured using Atomic Absorption Spectrometry (AAS, Perkin Elmer in LP2M) and by MC-ICP-MS during isotopic analyses (Thermofischer Sci. in ENS-Lyon, National CNRS-INSU Isotopic Platform), with a volume of 2.6 10^3^ cubic μm/cell.

### NHE1 activity measurement

Cells were loaded with BCECF/AM and excited at 490 and 450 nm. Images were recorded and treated as described in [Milosavljevic15]. Calibration was performed using 140 mM K^+^, 5 mM nigericin solutions ranging from pH 6.5 to 7.4.

### Li purification for isotopic analyses

After the experiments, collected cells and lysates were treated with concentrated and trace-cleaned acids, and Li extracted and purified in a clean laboratory for preparing them to isotope analyses. In brief, cells were dissolved using concentrated HNO_3_ and H_2_O_2_, followed by reverse *aqua regia*, in order to obtain a complete dissolution. All samples were then dried and dissolved in titrated ultrapure 1.0 M HCl, and centrifuged. The dissolution was considered as complete if no residue was visible at this step. Then, Li separation and purification were performed using 8 cm high AG50-X8 cation exchange resin columns, following a well-established procedure^19–21^. Each sample was passed through the cationic exchange resin columns twice, in order to fully purify the Li fractions from the sample matrix.

The solutions were dried and hot residues dissolved in 0.05M HNO_3_ for isotope analyses.

### Li isotope analyses

For all samples, cells and solutions, Li isotope analyses were performed using the Neptune *Plus* MC-ICP-MS (Multi Collector Ion Coupling Plasma Mass Spectrometry) available in the National CNRS-INSU Facilities platform at ENS-Lyon. The corresponding configuration use an Aridus II desolvating system, and specific high sensitivity cones. Thus, the sensitivity is about 1V/ppbLi, with a small, constant, and regularly monitored memory effect (typically less than 2% of the ^7^Li signal), necessary to measure low Li level biological materials. The standard bracketing technique is used along with a systematic blank correction before each sample and standard, in order to correct from the instrumental isotope fractionation. Several reference materials are run during each measurement session, typically Li7-N, Seawater and LiCl Sigma solution. When possible, analytical replicates were performed in order to verify the representative nature of the measured isotope ratio. Finally, full replicates including the whole protocol from the Li uptake by cells, dissolution, and Li purification and Li isotope analyses, in order to quantify the reproducibility of our methods. All corresponding uncertainties are reported in the Supplementary Tables.

### Electrophysiological measurements of Li transport

Membrane fractions from PS120 cells were prepared using mechanical disruption by the nitrogen decompression method (Parr cell disruption vessel). This step was followed by sucrose fractionation, solubilization in an appropriate non-ionic and non-denaturing detergent (Active Motif, USA), and incorporation into artificial lipid bilayers (Orbit mini, Nanion, Germany). Briefly, lipid bilayers are formed using 1,2-diphytanoyl-sn-glycero-3-phopsphocholine (DPHPC) lipid (10 mg/mL in octane), and membrane fractions were reincorporated by applying the appropriate amount of proteins to the upper chamber to elicit single channel activity. Single channel recordings were performed in a symmetrical solution (140 mM Li-acetate, pH7.4) to elicit only Li^+^ currents.

