## supplementary tables, figure and full mathematical model for "Biological fractionation of lithium isotopes by cellular Na^+^/H^+^ exchangers unravels fundamental transport mechanisms"

‡ Co-first authors

### SUPPLEMENTARY INFORMATION

### 1. Tables of results for Li transport

**Table S1.** Kinetic Li and Li isotopic active transport through membranes of NHE1 equipped cells. For all experiments, the external solution Li concentration was 15mM and its $\delta^7$Li value is 15.0‰ +/-0.3‰ ($2\sigma_n$, n=10). n is the number of full replicates (from experiment to isotope analysis).

| Cell line | T (°C) | Ext Li (mM) | Uptake duration (s) | Intracell Li (mM) | $2\sigma_n$ | $\delta^7$Li (‰) | $2\sigma_n$ | n |
|---|---|---|---|---|---|---|---|---|
| NHE-1 | 37 | 15 | 0 | 0.04 | | | | 1 |
| NHE-1 | 37 | 15 | 5 | 5.11 | 2.2 | 1.00 | 0.44 | 2 |
| NHE-1 | 37 | 15 | 10 | 10.49 | 5.3 | 0.83 | 0.30 | 2 |
| NHE-1 | 37 | 15 | 15 | 14.68 | 3.9 | 1.31 | 0.72 | 4 |
| NHE-1 | 37 | 15 | 30 | 24.22 | 2.8 | 1.57 | 0.19 | 3 |
| NHE-1 | 37 | 15 | 60 | 35.19 | 3.7 | 2.55 | 0.94 | 9 |
| NHE-1 | 37 | 15 | 180 | 66.17 | 29.3 | | | 1 |
| NHE-1 | 37 | 15 | 300 | 101.38 | 10.1 | 7.54 | 0.40 | 1 |
| NHE-1 | 37 | 15 | 900 | 108.81 | 10.9 | 12.64 | 0.40 | 1 |
| NHE-1 | 37 | 15 | 3600 | 88.52 | 21.9 | 13.24 | 1.73 | 3 |

**Table S2.** Kinetic Li passive transport through membrane of cells which do not express any NHE (PS120 or NHE-Null cell). For all experiments, the external solution Li concentration was 15mM.

| Cell line | T (°C) | Ext Li (mM) | Uptake duration (s) | Intracell Li (mM) | n |
|---|---|---|---|---|---|
| PS120 | 37 | 15 | 15 | 0.75 | 3 |
| PS120 | 37 | 15 | 30 | 0.77 | 3 |
| PS120 | 37 | 15 | 60 | 1.64 | 3 |
| PS120 | 37 | 15 | 180 | 4.03 | 3 |
| PS120 | 37 | 15 | 360 | 7.10 | 3 |
| PS120 | 37 | 15 | 600 | 9.61 | 3 |
| PS120 | 37 | 15 | 1200 | 14.3 | 3 |
| PS120 | 37 | 15 | 1800 | 15.5 | 3 |
| PS120 | 37 | 15 | 2400 | 19.6 | 3 |
| PS120 | 37 | 15 | 3600 | 24.7 | 3 |

**Table S3.** Dose Response of Li and Li isotopes active (NHE1) and passive (Cariporide inhibitor) transport through membrane of cells equipped with NHE1. For all experiments, the duration of the Li uptake was 60 s. The external solution $\delta^7$Li value is 15.0‰ +/-0.3‰ (2$\sigma_n$, n=10).

| Cell line | T (°C) | Ext Li (mM) | Intracell Li (mM) | $2\sigma_n$ | $\delta^7$Li (‰) | $2\sigma_n$ | n |
|---|---|---|---|---|---|---|---|
| NHE1 | 37 | 0 | 0.04 | 0.004 | | | 1 |
| NHE1 | 37 | 0.3 | 3.16 | 0.32 | 5.37 | 0.02 | 2 |
| NHE1 | 37 | 1 | 6.39 | 4.9 | 3.86 | 1.37 | 4 |
| CARIPORIDE* | 37 | 1 | 0.10 | 0.004 | 8.89 | 0.48 | 2 |
| NHE1 | 37 | 3 | 11.62 | 8.0 | 3.92 | 1.34 | 5 |
| CARIPORIDE* | 37 | 3 | 0.19 | 0.04 | 8.26 | 1.49 | 2 |
| NHE1 | 37 | 15 | 35.19 | 3.7 | 2.55 | 0.94 | 9 |
| NHE1 | 20$^+$ | 15 | 35.94 | 11.4 | 2.71 | 0.24 | 1 |
| NHE1 | 37 | 30 | 30.44 | 23.0 | 1.88 | 0.40 | 4 |
| CARIPORIDE* | 37 | 30 | 1.51 | 0.33 | 6.72 | 0.17 | 2 |
| NHE1 | 37 | 60 | 11.85 | 0.59 | 1.43 | 0.25 | 2 |
| NHE1 | 37 | 90 | 12.57 | 0.10 | 1.09 | 0.23 | 2 |
| NHE1 | 37 | 120 | 1.81 | 0.18 | 1.61 | 0.02 | 1 |

*Cariporide inhibits the Li active transport by NHEs, as evidenced by the low intracellular Li measured at the end of the experiments.
$^+$This experience has been performed at 20°C for comparison

**Table S4**. Li and Li isotopic active transport by NHE1 equipped cells when the external solution contains variable amounts of Na. For all experiments, the duration of the Li and Na uptake was 60 s. For all experiments, the external solution Li concentration is 15mM and its $\delta^7$Li value is 15.0‰ +/-0.3‰ (2$\sigma_n$, n=10).

| Ext Na (mM) | T (°C) | Ext Li (mM) | Intracell Li (mM) | $2\sigma_n$ | $\delta^7$Li (‰) | $2\sigma_n$ | n |
|---|---|---|---|---|---|---|---|
| 0 | 37 | 15 | 35.19 | 3.7 | 2.55 | 0.94 | 9 |
| 10 | 37 | 15 | 32.26 | 1.71 | 2.51 | 0.20 | 3 |
| 30 | 37 | 15 | 28.90 | 2.59 | 2.38 | 0.36 | 3 |
| 60 | 37 | 15 | 23.23 | 5.25 | 3.31 | 0.25 | 3 |
| 105 | 37 | 15 | 15.35 | 1.53 | 2.76 | 0.02 | 1 |

**Table S5**. Li and Li isotopic active transport by NHE1 equipped cells when the internal solution contains variable amounts of protons (pH different). For all experiments, the duration of the Li uptake was 60 s. For all experiments, the external solution Li concentration is 15mM and its $\delta^7$Li value is 15.0‰ +/-0.3‰ (2$\sigma_n$, n=10).

| Intracell pH* | T (°C) | Ext Li (mM) | Intracell Li (mM) | $2\sigma_n$ | $\delta^7$Li (‰) | $2\sigma_n$ | n |
|---|---|---|---|---|---|---|---|
| 7.30 | 37 | 15 | 1.10 | 0.05 | 16.7 | 0.9 | 2 |
| 6.85 | 37 | 15 | 2.79 | 0.65 | 7.6 | 2.0 | 3 |
| 6.51 | 37 | 15 | 17.66 | 2.80 | 2.7 | 0.1 | 3 |
| 6.30 | 37 | 15 | 26.25 | 4.57 | 2.1 | 0.4 | 3 |
| 6.24 | 37 | 15 | 34.52 | 2.66 | 2.0 | 0.1 | 2 |
| 6.13 | 37 | 15 | 35.19 | 3.7 | 2.5 | 0.9 | 9 |

*intracellular pH were reached using various $NH_4Cl$ uptake times, and was estimated using the calibration described in the Method section.

### 2. Figure S1: $Li^+$ currents in lipid bilayers

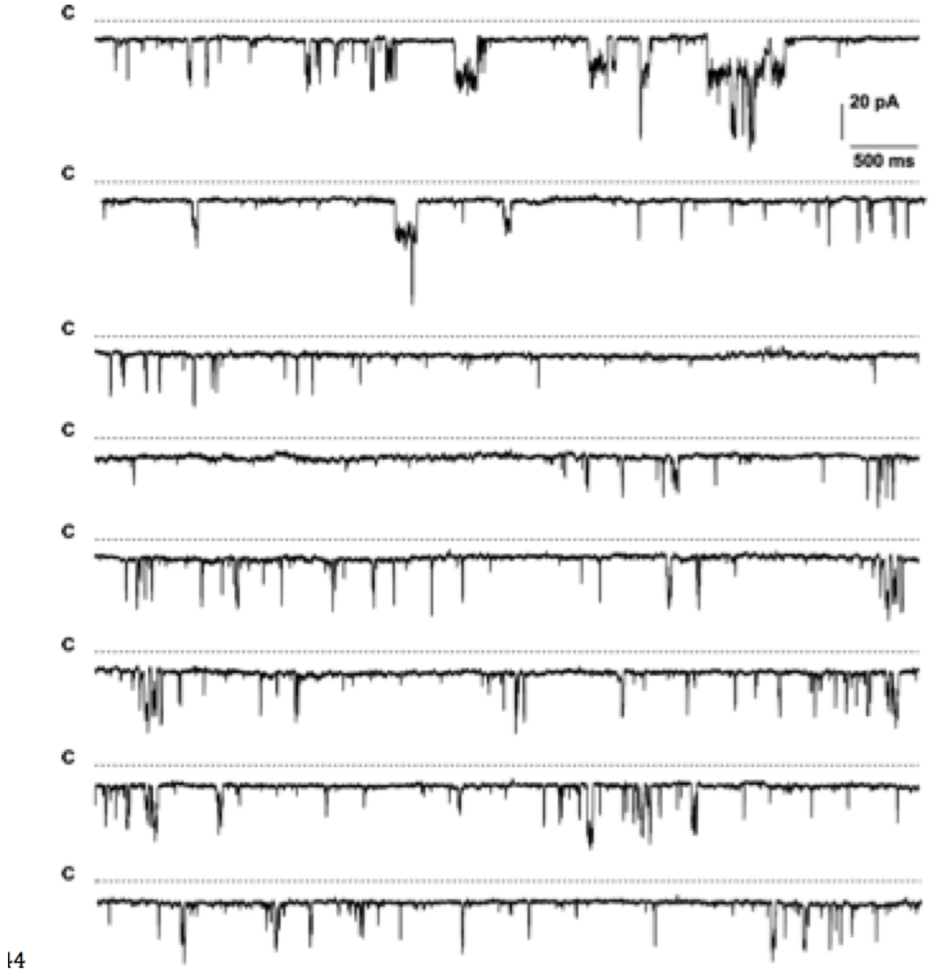


NHE-1-expressing PS120 membrane fragments were incorporated into lipid bilayers and current recordings were performed at various continuous potentials. We show a representative current gating from a closed state (C) elicited by application of a potential of -100 mV. Single channel opening events appear as downward deviations of the baseline, which correspond to Li currents in the range of the scale (20 pA) and of various frequencies and durations (from several ms to >100ms).

### 3. Supplementary Kinetic Model Construction

#### 3.I . CELLULAR SYSTEM WITH ISOTOPIC FRACTIONATION

We consider a cell prepared according to the protocol described in the main manuscript, and we deal with a cell model as described by Figure 1A. The cytosolic concentrations and variables are named without any specific index, and the outer concentrations and variables are named with a '$out$' index. Hence, the variables of interest are:

- $\Lambda$ as the total lithium concentration,
- $E_m$ as the membrane potential (see §6.III for its derivation),
- $[{}^6Li^+]_{out} = \varepsilon_6\Lambda_{out}$, where $\varepsilon_6 \simeq 0.076$ is the natural abundance of ${}^6Li^+$ in the outer lithium solution,
- $[{}^7Li^+]_{out} = \varepsilon_7\Lambda_{out}$, where $\varepsilon_7 \simeq 0.924$ is the natural abundance of ${}^7Li^+$ in the outer lithium solution.

The Table I sums up the different initial values that we need to start a kinetic model.

| Initial Value | Outer Medium | Cytosol |
|---|---|---|
| $[Li^+]$ | $\Lambda_{out}$ | $\Lambda = 0$ |
| $[{}^6Li^+]$ | $\varepsilon_6\Lambda_{out}$ | 0 |
| $[{}^7Li^+]$ | $\varepsilon_7\Lambda_{out}$ | 0 |
| $[K^+]$ | 10 mM | 140 mM |
| $[Cl^-]$ | 140 mM | 20 mM |
| $\delta^7Li^+$ | 14.57 | N/A |
| $E_m$ | – | -40 mV |

TABLE I: Summary of values accounting for the initial cell state.

#### 3. II. TRANSPORTING ISOTOPES BY AN ACTIVATED PROCESS

##### A. Description

As referred to in the main manuscript, a recent study showed how one monovalent cation is driven across one NHE monomer with the occurence of a local flip, leading the cation from a nanoscopic groove close to the outer medium (*ie* the ”outer groove”, outlined in blue on the left of Figure 1B) to a nanoscopic groove close to the cytosol (*ie* the ”cytosolic groove”, outlined in orange on the right of Figure 1B). We assume that this molecular flipping mechanism is the rate determining step of the lithium translocation, and the free energy accompanying this transformation is sketched on Figure 1B, where all the relevant free energies are reported. We classically consider that there is no significant energy well near the unbounded free energy maximum.

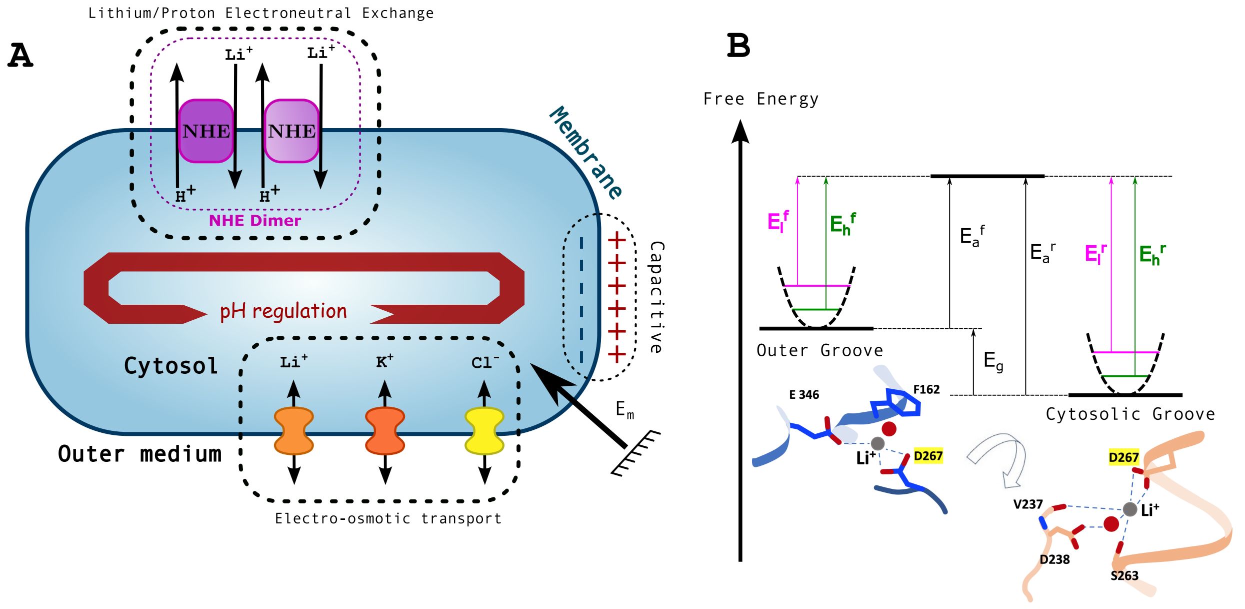


FIG. 1: **A**. *Representation of the different kinetic elements.* **B**. *Schematic translocation from the outer groove to a more stable cytosolic groove within one NHE monomer. The absolute free energy gain is $E_g$, while the absolute activation energies are shown as $E_a$. A light element ( $^6Li^+$ in magenta) starts from a quasi-harmonic state (see § 6.II B) and has a forward activation energy $E_l^f$ and a backward activation free energy $E_l^b$. A heavy element ( $^7Li^+$ in green) starts from a quasi-harmomic state and has a forward activation energy $E_h^f$ and a backward activation energy $E_h^b$.*

#### B. Harmonic Approximations and Reduced Masses

We assume here that the energy minima are approximated by two harmonic potentials with a stiffness of $k_r$ for the ”reactants” (outer groove) and $k_p$ for the

”products” (cytosolic groove). Accordingly, the fundamental energy levels $E_r$ and $E_p$ are respectively :

$$\begin{cases} E_r(\mu) &= \frac{1}{2}\hbar\sqrt{\frac{k_r}{\mu}} \\ E_p(\mu) &= \frac{1}{2}\hbar\sqrt{\frac{k_p}{\mu}} - E_g, \end{cases} \tag{1}$$

where $\mu$ is the reduced mass of the interacting particules and $\hbar$ is the reduced Plank’s constant. Using this result and the notation of § 6.II A and Figure 1, we shall now derive the differences in thermodynamics and kinetics, depending on the reduced masses.

Moreover, the reduced mass is always close to the mass of the lightest particle that is involved in the chemical transformation. Hence we expect $\mu_l$ to be the mass of $^6Li^+$ and $\mu_h$ to be the mass of $^7Li^+$.

#### C. Isotopic Thermodynamics

For a given reduced mass $\mu$, the total difference of energy between the states can be described as:

$$\Delta E(\mu) = E_p(\mu) - E_r(\mu) = -E_g + \frac{\hbar}{2\sqrt{\mu}}\left(\sqrt{k_p} - \sqrt{k_r}\right). \tag{2}$$

If we now use $\mu_l$ for the lighter system and $\mu_h$ for the heavier system, the difference of energy differences is:

$$\underbrace{\Delta E(\mu_l) - \Delta E(\mu_h)}_{\Delta\Delta E} = \frac{\hbar}{2}\left(\sqrt{k_p} - \sqrt{k_r}\right)\underbrace{\left(\frac{1}{\sqrt{\mu_l}} - \frac{1}{\sqrt{\mu_h}}\right)}_{>0}. \tag{3}$$

- If the species senses the same environment $(k_r \simeq k_p)$, then the energy difference is the same and there is no influence of the mass on the thermodynamics.
- Otherwise, a slight change in composition may occur.
  - If the outer groove potential well is less stiff than the extracellular one $(k_p < k_r)$, then there will be an excess of lighter products $(\Delta\Delta E < 0)$.
  - Conversely, if the cytosolic groove potential well is stiffer than the extracellular one. $(k_r < k_p)$, then there will be an excess of heavier products $(\Delta\Delta E > 0)$.

#### D. Isotopic Kinetics

Using the notations of § 6.II A and § 6.II C, the forward and backward activation energies are respectively:

$$\begin{cases} E_a^f(\mu) = E_a - E_r(\mu) \\ E_a^b(\mu) = E_a - E_p(\mu) \end{cases} \tag{4}$$

Accordingly, the apparent first order rate constants are respectively:

$$\begin{cases} k_a^f(\mu) = k_0^f e^{\left[\frac{E_r(\mu)}{k_B T} - \frac{E_a}{k_B T}\right]} \\ \\ k_a^b(\mu) = k_0^b e^{\left[\frac{E_p(\mu)}{k_B T} - \frac{E_a}{k_B T}\right]} . \end{cases} \tag{5}$$

Let us now evaluate the ratio of the lighter forward rate over the heavier forward rate:

$$\begin{cases} \frac{k_a^f(\mu_l)}{k_a^f(\mu_h)} & = \exp\left[\frac{E_r(\mu_l) - E_r(\mu_h)}{k_B T}\right] \\ \\ & = \exp\left[\frac{\hbar\sqrt{k_r}}{2k_B T}\underbrace{\left(\frac{1}{\sqrt{\mu_l}} - \frac{1}{\sqrt{\mu_h}}\right)}_{>0}\right] \\ \\ & = \kappa_f > 1. \end{cases} \tag{6}$$

Similarly, the ratio of the lighter backward rate over the heavier backward rate can be written:

$$\frac{k_a^b(\mu_l)}{k_a^b(\mu_h)} = \exp\left[\frac{\hbar\sqrt{k_p}}{2k_B T}\underbrace{\left(\frac{1}{\sqrt{\mu_l}} - \frac{1}{\sqrt{\mu_h}}\right)}_{>0}\right] = \kappa_b > 1. \tag{7}$$

#### E. Conclusions

When two isotopes undergo the same transformation in a potential energy landscape, some differences occur in both thermodynamics and kinetics. Actually, the lighter isotope preferentially stays in the less constraint spaces (a.k.a the heavier isotope preferentially stays in the most constraint spaces), and always has a faster way to reach its equilibria.

Moreover, if two isotopes undergo a set of transformations leading to different potential wells with stiffness $k_{p,j\in[1,n]}$, then the apparent initial speedup $\kappa_n^f$ is expressed by:

$$\begin{aligned}
\kappa_{f,n} &= \exp\left[\frac{\hbar}{2k_BT}\left(\frac{1}{\sqrt{\mu_l}}-\frac{1}{\sqrt{\mu_h}}\right)\left(\sum_{j=1}^{n}\sqrt{k_{p,j}}\right)\right] \\
&= \exp\left[\frac{\hbar}{2k_BT}\left(\frac{1}{\sqrt{\mu_l}}-\frac{1}{\sqrt{\mu_h}}\right)n\frac{1}{n}\quad\sum_{j=1}^{n}\sqrt{k_{p,j}}\right)\right] \\
&= \exp\left[\frac{\hbar}{2k_BT}\left(\frac{1}{\sqrt{\mu_l}}-\frac{1}{\sqrt{\mu_h}}\right)\langle k_p\rangle\right]^n \\
&= \left\langle\kappa_f\right\rangle^n
\end{aligned} \tag{8}$$

### 3 .III. ELECTRO-OSMOTIC TRANSPORT

#### A. Leaks arising from the electro-osmotic gradients

The net charge difference between the cytoplasm and the outer medium gives birth to a cell membrane polarisation $E_m$. Consequently, each ion leaks through membrane channels according to both its specific permeability and the Goldman-Hodgkin-Katz (GHK) flux equations. We sum up here the corresponding intakes for an anion $A^-$ or a cation $Q^+$ whilst relying on the formalism of [Bouret14], where $k_A$ and $k_Q$ are their respective full-cell permeabilities:

$$\left\{\begin{aligned}
\zeta &= \frac{\mathcal{F}E_m}{RT} && \text{(reduced potential, } \mathcal{F} = \text{Faraday constant)} \\
\Psi(\zeta) &= \frac{\zeta}{1-e^{-\zeta}} \simeq 1+\frac{1}{2}\zeta+\frac{1}{12}\zeta^2+\ldots && \text{(GHK current weight function)} \\
\partial_t[Q^+]|_{leak} &= k_Q\Psi(\zeta)\left([Q^+]_{out}e^{-\zeta}-[Q^+]\right) && \text{(full cell leak for a cation } Q^+\text{)} \\
\partial_t[A^-]|_{leak} &= k_A\Psi(\zeta)\left([A^-]_{out}-[A^-]e^{-\zeta}\right) && \text{(full cell leak for an anion } A^-\text{)}
\end{aligned}\right. \tag{9}$$

The net balance of all the species $X_i$ with a charge $z_i$ leads to the electric equation, depending on the full-cell membrane capacitance $C_m$:

$$C_m \partial_t E_m = \mathcal{F} \sum_i z_i \partial_t [X_i]|_{leak}, \tag{10}$$

and we define the electric sensitivity $\gamma$ as:

$$\gamma = \frac{\mathcal{F}}{C_m}. \tag{11}$$

#### B. Application to NHE-null (PS120) cells

##### 1. *Corresponding differential system*

As formerly shown [Bouret14] and as there is no present $Na^+$ in these sets of experiments, the main ions that account for the potential evolution are $K^+$ and $Cl^-$, with $k_{Cl} \approx 0.1 k_K$. We obviously take into account the lithium contributions with $k_7$ for $^7Li^+$ and $k_6$ for $^6Li^+$. In water [Richter06], the diffusion coefficient $D_{^6Li^+}$ of $^6Li^+$ is greater than the one $D_{^7Li^+}$ for $^7Li^+$ by a ratio:

$$\frac{D_{^7Li^+}}{D_{^6Li^+}} = 0.99772 \pm 0.00026, \tag{12}$$

which is equivalent to a diffusion speed-up of $^6Li^+$ compared to $^7Li^+$ of:

$$\sigma = \frac{D_{^6Li^+}}{D_{^7Li^+}} = 1.00229 \pm 2 \cdot 10^{-4} \tag{13}$$

Since the transport of species through ion channels is considered as a diffusive process, we define the ”electro-osmotic acceleration”:

$$\sigma' = \frac{k_6}{k_7}. \tag{14}$$

Finally, the NHE-null differential system is:

$$\begin{cases} \partial_t[\,^6Li^+] &= k_6 \Psi(\zeta)\left([\,^6Li^+]_{out} e^{-\zeta} - [\,^6Li^+]\right) = \partial_t[\,^6Li^+]|_{leak} \\ \partial_t[\,^7Li^+] &= k_7 \Psi(\zeta)\left([\,^7Li^+]_{out} e^{-\zeta} - [\,^7Li^+]\right) = \partial_t[\,^7Li^+]|_{leak} \\ \partial_t[K^+] &= k_K \Psi(\zeta)\left([K^+]_{out} e^{-\zeta} - [K^+]\right) \\ \partial_t[Cl^-] &= k_{Cl} \Psi(\zeta)\left([Cl^-]_{out} - [Cl^-] e^{-\zeta}\right) \\ \partial_t E_m &= \gamma\left(\partial_t[\,^6Li^+]|_{leak} + \partial_t[\,^7Li^+]|_{leak} + \partial_t[K^+] - \partial_t[Cl^-]\right) \end{cases} \tag{15}$$

### 2. *Estimation of the constants*

We designed a $C^{++}$ software to solve the differential problem (using a Dormand-Price algorithm), coupled with a Levenberg–Marquardt method to adjust the different parameters for the following constraints:

| Parameter | Value | Origin |
|---|---|---|
| $\Lambda_{out}$ | 15 mM | imposed concentration |
| $\delta^7 Li^+\|_{t=60s}$ | 10.8 | average measured value |
| $k_{Cl}$ | $0.1 k_K$ | see electrophysiology in [Bouret14] |

We obtain the following estimations:

$$\begin{cases} \gamma = (5.21 \pm 0.38) & \mathrm{V.M^{-1}} \\ k_K = (0.058 \pm 0.001) & \cdot 10^{-3}\ \mathrm{s^{-1}} \\ k_7 = (0.992 \pm 0.04) & \cdot 10^{-3}\ \mathrm{s^{-1}} \end{cases} \quad (16)$$

and we deduced from the computation that the ratio of the lithium permeabilities is:

$$\sigma' = \frac{k_6}{k_7} = 1.003785 \pm 3 \cdot 10^{-6}, \quad \frac{1}{\sigma'} = 0.996229 \pm 3 \cdot 10^{-6}, \quad (17)$$

that consistently yields a computed value of $\delta^7 Li^+ = 10.8 \pm 0.9$ at one minute of uptake, in accordance with the required constraints. The resulting adjusted curve is shown in the main manuscript on Figure 3a.

### 3. *Conclusion from NHE-null measurements*

The kinetic data are fully coherent with an electro-osmotic regulation of a NHE-null cell undergoing our experimental protocol. The value of the greater selectivity for $^6Li^+$ through the ionic channels agrees with a diffusive process, as hypothesised by Goldman-Hodgkin-Katz, and this is the first time to our knowledge that a measure of this physiological constant is provided.

### 1. I V. COOPERATIVE LITHIUM TRANSPORT BY NHE

#### A. Model for a dimer

##### 1. *Combinatorics of membrane species*

Let's name the bare (*ie* without any linked ion) monomer $\mathcal{E}$. Then the bare dimer $2\mathcal{E}$ offers a side $a$ and a side $b$, with two binding sites on each side. We assume that each of these sites independently accepts an ion $X_1,\ldots,X_N$, to form a pre-equilibrated species as follow:

$$\forall(i,j,k,l)\in[1:N]^4,\;\; X_{(i,a)}+X_{(k,a)}+2\mathcal{E}+X_{(j,b)}+X_{(l,n)} \rightleftharpoons \left\{\begin{array}{ccc} X_{(i,a)} & \cdot\mathcal{E}\cdot & X_{(j,b)} \\ X_{(k,a)} & \cdot\mathcal{E}\cdot & X_{(l,n)} \end{array}\right\}, \tag{18}$$

that is described as the dimeric association of two "$X\cdot\mathcal{E}\cdot X$" protomers.

For each set of $N$ concentrations on side $a$, $N$ concentrations on side $b$, and the total $[\mathcal{E}]$ we derive the $N^4$ pre-equilibria observable constants:

$$\forall(i,j,k,l)\in[1:N]^4,\;\; K_{ijkl}=\frac{\left[\left\{\begin{array}{ccc} X_{(i,a)} & \cdot\mathcal{E}\cdot & X_{(j,b)} \\ X_{(k,a)} & \cdot\mathcal{E}\cdot & X_{(l,n)} \end{array}\right\}\right]}{[\mathcal{E}]^2[X_{(i,a)}][X_{(k,a)}][X_{(j,b)}][X_{(l,n)}]}. \tag{19}$$

Here, for the two lithium isotopes and $[H^+]$, we deal with $N=3$ and $81$ species. From now on, we note $\vec{X}_a$ the vector of concentrations on side $a$, and $\vec{X}_b$ the vector of concentrations on side $b$. Namely, we obtain:

$$\vec{X}_{a/b}=\begin{bmatrix}[H^+]_{a/b}\\ [\,^6Li^+]_{a/b}\\ [\,^7Li^+]_{a/b}\end{bmatrix} \tag{20}$$

##### 2. *Free NHE computation*

Let $\mathcal{E}_0$ be the total concentration of NHE. The enzyme conservation writes:

$$\mathcal{E}_0=[\mathcal{E}]+2[\mathcal{E}]^2\underbrace{\sum_{i,j,k,l}K_{ijkl}[X_{(i,a)}][X_{(k,a)}][X_{(j,b)}][X_{(l,b)}]\Big)}_{\Xi(\vec{X}_a,\vec{X}_b)}, \tag{21}$$

leading to

$$[\mathcal{E}]=\mathcal{E}_0\frac{2}{1+\sqrt{1+8\mathcal{E}_0\Xi(\vec{X}_a,\vec{X}_b)}}=\frac{\mathcal{E}_0}{W(\vec{X}_a,\vec{X}_b)} \tag{22}$$

#### 3. Conditional exchange (flip) rates

Each formed dimer formally flips its content with $2 \times 2$ rates, as expected from Eq.(18). We use a two sets of rates $U_{ijkl}$ for the "upper" protomer and $V_{ijkl}$ for the "lower" protomer to obtain:

$$\left\{ \begin{array}{l} \left\{ \begin{array}{ccc} X_{(i,a)} & \cdot\mathcal{E}\cdot & X_{(j,b)} \\ X_{(k,a)} & \cdot\mathcal{E}\cdot & X_{(l,n)} \end{array} \right\} \underset{U_{jikl}}{\overset{U_{ijkl}}{\rightleftharpoons}} \left\{ \begin{array}{ccc} X_{(j,a)} & \cdot\mathcal{E}\cdot & X_{(i,b)} \\ X_{(k,a)} & \cdot\mathcal{E}\cdot & X_{(l,n)} \end{array} \right\} \\ \\ \left\{ \begin{array}{ccc} X_{(i,a)} & \cdot\mathcal{E}\cdot & X_{(j,b)} \\ X_{(k,a)} & \cdot\mathcal{E}\cdot & X_{(l,n)} \end{array} \right\} \underset{V_{ijlk}}{\overset{V_{ijkl}}{\rightleftharpoons}} \left\{ \begin{array}{ccc} X_{(j,a)} & \cdot\mathcal{E}\cdot & X_{(i,b)} \\ X_{(l,a)} & \cdot\mathcal{E}\cdot & X_{(k,n)} \end{array} \right\} \end{array} \right. \tag{23}$$

Using the pre-equilibria from Eq.(19) and knowing that $[E]$ is expressed by Eq.(22), the creation rate of each species is deduced as the following example:

$$\begin{aligned} \frac{1}{[\mathcal{E}]^2}\partial_t [X_{(\mathbf{i},a)}] \; = \; & -[X_{(\mathbf{i},a)}] \sum_{jkl} \left(U_{\mathbf{i}jkl} K_{\mathbf{i}jkl} + V_{kl\mathbf{i}j} K_{kl\mathbf{i}j}\right) [X_{(j,b)}][X_{(k,a)}][X_{(l,b)}] \Big) \\ & +[X_{(\mathbf{i},b)}] \sum_{jkl} \left(U_{j\mathbf{i}kl} K_{j\mathbf{i}kl} + V_{klj\mathbf{i}} K_{klj\mathbf{i}}\right) [X_{(j,b)}][X_{(k,a)}][X_{(l,b)}] \Big) . \end{aligned} \tag{24}$$

We employ this formula to write the differential evolution of each chosen species.

#### 4. Initial isotopic separation and dose-response

We write the initial flip rates for the lithium isotopes:

- using no index for intracellular species and a *'out'* index for extracellular species (as in §6.I),
- using Eq.(24) to express individual rates,
- and using a '0' index for the initial time,

and we end up with (see §6.I for the definition of $\varepsilon_{6/7}$):

$$\left\{ \begin{array}{rcl} w_{pq} & = & U_{HpHq} K_{HpHq} + V_{HqHp} K_{HqHp}, \; p \in [6:7], \; q \in [6:7]. \\ \\ \left( \dfrac{[\,^7Li^+]}{[\,^6Li^+]} \right)_0 & = & \left( \dfrac{[\,^7Li^+]_{out}}{[\,^6Li^+]_{out}} \right) \dfrac{k_7 \Psi(\zeta_0) e^{-\zeta_0} + [H^+]_0^2 [\mathcal{E}]_0^2 \left(w_{76}\varepsilon_6 + w_{77}\varepsilon_7\right) \Lambda_{out}}{k_6 \Psi(\zeta_0) e^{-\zeta_0} + [H^+]_0^2 [\mathcal{E}]_0^2 \left(w_{66}\varepsilon_6 + w_{67}\varepsilon_7\right) \Lambda_{out}} \end{array} \right. \tag{25}$$

We clarify this expression by defining:

$$\left\{\begin{array}{rcl}\kappa' & = & \dfrac{w_{76}\varepsilon_6 + w_{77}\varepsilon_7}{w_{66}\varepsilon_6 + w_{67}\varepsilon_7} \\ \\ \tan^2\omega & = & \dfrac{(w_{76}\varepsilon_6 + w_{77}\varepsilon_7)\,[H^+]_0^2[\mathcal{E}]_0^2\Lambda_{out}}{\Psi(\zeta_0)e^{-\zeta_0}},\end{array}\right. \tag{26}$$

and we consequently obtain:

$$\left(\frac{[\,^7Li^+]}{[\,^6Li^+]}\right)_0 = \left(\frac{[\,^7Li^+]_{out}}{[\,^6Li^+]_{out}}\right)\frac{1}{\sigma'\cos^2\omega + \kappa'\sin^2\omega} = \left(\frac{[\,^7Li^+]_{out}}{[\,^6Li^+]_{out}}\right)\frac{1+\tan^2\omega}{\sigma' + \kappa'\tan^2\omega}. \tag{27}$$

We remind the following expression:

$$\delta\,^7Li_0^+ = 10^3\left[(1+10^{-3}\delta\,^7Li_{out}^+)\frac{1+\tan^2\omega}{\sigma'+\kappa'\tan^2\omega} - 1\right], \tag{28}$$

and we want to find out how the initial fractionation depends on the total extracellular lithium concentration $\Lambda_{out}$, which is equivalent to study the variations of:

$$\Xi = \frac{1+\tan^2\omega}{\sigma'+\kappa'\tan^2\omega}. \tag{29}$$

Since $\tan\omega^2 > 0$ and the experimental initial concentration ratio is smaller than the external one, we deduce that $\kappa' > \sigma'$. Accordingly, we observe that $\Xi$ is a *decreasing* function with respect to $\tan^2\omega$, which is proportional to $[\mathcal{E}]_0^2\Lambda_{out}$:

- this expression starts from zero, meaning that the first-order in $\Lambda_{out}$, GHK pathway always imposes of fractionation related to $\sigma'$,
- and this expression returns to zero for high $\Lambda_{out}$, since there is a saturation in the NHE kinetics, while the GHK pathways is always proportional to $\Lambda_{out}$,
- thus, since $[\mathcal{E}]_0^2\Lambda_{out}$ is positive and starts from zero and returns to zero, it is *increasing* on an interval starting from zero.

Consequently, we directly deduce that, due to the very dimeric structure of NHE:

- there exists *an optimal total outer lithium concentration that we name $\Lambda_{opt}$ that produces a maximum initial fractionation,*
- there is an *increasing fractionation* with respect to an *increasing total outer lithium concentration* ranging from $0$ to $\Lambda_{opt}$ (synergic dose-response),

- and that the experimental data shows (on Figure 4e of the main article) that the value of $\Lambda_{opt}$ for the studied cells is about $100\text{mM}$, and therefore allows the observation of the dose-response as described above.

#### 5. *Dose-response morphology*

Since $\Xi$ has a very complex expression, we choose a simplified set of parameters:

- $\varepsilon_6 \ll \varepsilon_7$,
- $[H^+]_{out} \ll \Lambda_{out}$,
- $\forall i,j,k,l,\ K_{ijkl} \simeq K_0$ [see §(6.IV C 2) for a formal justification].

By a continuity argument, the real behaviour of $\Xi$ will have the same traits than this simplified version that we call $\tilde{\Xi}$. We obtain a transformed expression with the help of a scaling factor $\chi_7$ and a *reduced concentration* $\lambda$:

$$\begin{cases} \chi_7 = \dfrac{w_{77}}{\Psi(\zeta_0)e^{-\zeta_0}}\sqrt{\dfrac{2\mathcal{E}_0^3}{K_0}} \\ \\ \lambda = (2^{\frac{3}{2}}[H^+]_0\sqrt{\mathcal{E}_0 K_0})\Lambda_{out} \\ \\ \tilde{\Xi} = \dfrac{1}{\sigma'}\dfrac{\left[1+\sqrt{1+\lambda^2}\right]^2+\chi_7\lambda}{\left[1+\sqrt{1+\lambda^2}\right]^2+\dfrac{\kappa'}{\sigma'}\chi_7\lambda} \end{cases} \tag{30}$$

The *reduced concentration* $\lambda$ is a non-trivial combination of all the components acting on the fractionation efficiency, and shows in particular a proportionality to $\Lambda_{out}$ and the initial acidity $[H^+]_0$. Finally, we experimentally measured $\delta\,^7Li^+|_{t=60s}$, which is expected to be lower that $\delta\,^7Li_0^+$ due to the favoured entry of $^6Li^+$. That is why we propose the following expression:

$$\delta\,^7Li^+|_{t=60s} \propto 10^3\left[(1+10^{-3}\delta\,^7Li_{out}^+)\tilde{\Xi}-1\right]. \tag{31}$$

A simple fit procedure yields the following values, using $\kappa' \simeq 1.0148$ (see further Eq.(41)):

$$\begin{cases} (2^{\frac{3}{2}}[H^+]_0\sqrt{\mathcal{E}_0 K_0}) \simeq (46 \pm 5\text{ mM})^{-1} \\ \chi_7 \simeq 12 \pm 3 \\ \delta\,^7Li^+|_{t=60s} \simeq (0.45 \pm 0.05)\delta\,^7Li_0^+ \end{cases} \tag{32}$$

Obviously, these numbers are very rough and almost heuristic estimations, but we retrieve the predicted behaviour, as shown on Figure 4e with the adjusted curve.

##### 6. Conclusion of the role of NHE

We observe the following points:

- *(consistency)* without NHE, equivalent to $\omega = 0$ in Eq.(27), we retrieve the passive description, as seen in Eq.(17),
- *(cooperativity)* but with NHE and its cooperative (dimeric) action, an increasing fractionation is observed for an experimentally extended range of total lithium concentration,
- *(novelty)* which is of fundamental importance, since the same approach with the assumption that only a monomer exchanges the ions classically leads to a simple saturation function for the initial intake rate with respect to time, and no such dose-response could be predicted.

#### B. pH recovery

One the one hand, the extracellular pH is fixed by the experiment. But on the other hand, since the kinetic scheme depends on the intracellular $[H^+]$, other measurements were performed to monitor the pH rise after the initial acidification, and are reported on Figure 2 for different total external lithium concentrations. Since we do not include in this model the full homeostasis, we take $[H^+]$ as an experimental dependent input, and it appears that, with a good approximation, $[H^+]$ is a function of time $t$, with parameters $[H^+]_0$, $[H^+]_\infty$, and a recovery time $t_h$ such that:

$$[H^+] \simeq [H^+]_0 + \left([H^+]_\infty - [H^+]_0\right) \frac{t}{t+t_h} = [H^+]_\infty + \left([H^+]_0 - [H^+]_\infty\right) \frac{t_h}{t+t_h}. \quad (33)$$

We use this equation to fit the pH recovery data as indicated on Figure 2. It is important to remember that $[H^+]_\infty$ depends on $\Lambda_{out}$. From the measurements, we use an heuristic expression for $[H^+]$ dynamics depending on $\Lambda_{out}$:

$$\left\{\begin{array}{rcl} t_h & = & \dfrac{A_h}{\mathrm{erf}\,(B_h\Lambda_{out})} \\ A_h & = & 23.95 \pm 1.42 \text{ s} \\ B_h & = & 0.0318 \pm 0.004 \text{ mM}^{-1} \end{array}\right. \tag{34}$$

and

$$\left\{\begin{array}{rcl} \mathrm{pH}_0 & = & 5.94 \pm 0.1 \\ \mathrm{pH}_\infty & = & \mathrm{pH}_0 + (7.40 - \mathrm{pH}_0)\,\dfrac{\Lambda_{out}}{\Lambda_h + \Lambda_{out}} \\ \Lambda_h & = & 15.11 \pm 1.5 \text{ mM} \end{array}\right. \tag{35}$$

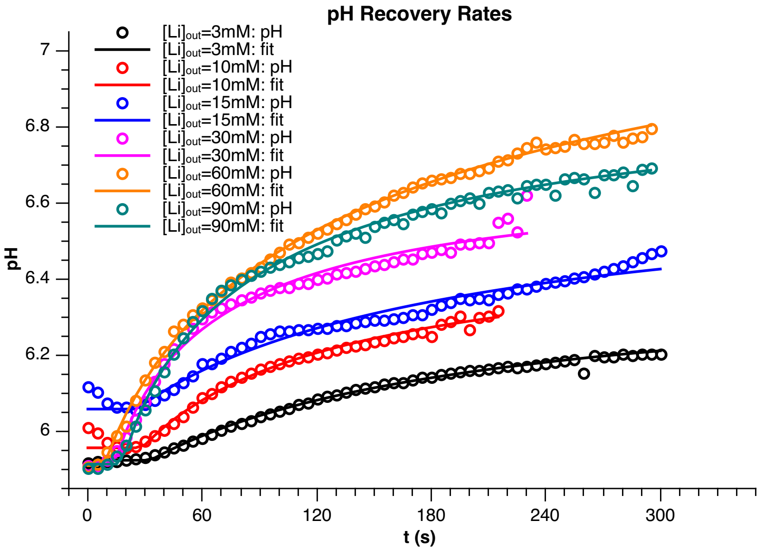


FIG. 2: *Adjustment (lines, according to Eq.* (33)*) of apparent experimental protons intake (open circles), for various total external lithium concentrations.*

#### C. Curves Adjustment

##### 1. *Kinetic Reduction*

We are left with 81 possible species. Let's see what are their contributions based on experimental and symmetry arguments.

- We observe that $\mathbf{9}$ symmetric species are ”futile”: their flips produce no concentration change and they act only as inhibitors:

$$\begin{array}{|c|c|c|} \hline \left\{\begin{array}{ccc} H^+ & \cdot\mathcal{E}\cdot & H^+ \\ H^+ & \cdot\mathcal{E}\cdot & H^+ \end{array}\right\} & \left\{\begin{array}{ccc} H^+ & \cdot\mathcal{E}\cdot & H^+ \\ {}^6Li^+ & \cdot\mathcal{E}\cdot & {}^6Li^+ \end{array}\right\} & \left\{\begin{array}{ccc} H^+ & \cdot\mathcal{E}\cdot & H^+ \\ {}^7Li^+ & \cdot\mathcal{E}\cdot & {}^7Li^+ \end{array}\right\} \\ \hline \left\{\begin{array}{ccc} {}^6Li^+ & \cdot\mathcal{E}\cdot & {}^6Li^+ \\ H^+ & \cdot\mathcal{E}\cdot & H^+ \end{array}\right\} & \left\{\begin{array}{ccc} {}^6Li^+ & \cdot\mathcal{E}\cdot & {}^6Li^+ \\ {}^6Li^+ & \cdot\mathcal{E}\cdot & {}^6Li^+ \end{array}\right\} & \left\{\begin{array}{ccc} {}^6Li^+ & \cdot\mathcal{E}\cdot & {}^6Li^+ \\ {}^7Li^+ & \cdot\mathcal{E}\cdot & {}^7Li^+ \end{array}\right\} \\ \hline \left\{\begin{array}{ccc} {}^7Li^+ & \cdot\mathcal{E}\cdot & {}^7Li^+ \\ H^+ & \cdot\mathcal{E}\cdot & H^+ \end{array}\right\} & \left\{\begin{array}{ccc} {}^7Li^+ & \cdot\mathcal{E}\cdot & {}^7Li^+ \\ {}^6Li^+ & \cdot\mathcal{E}\cdot & {}^6Li^+ \end{array}\right\} & \left\{\begin{array}{ccc} {}^7Li^+ & \cdot\mathcal{E}\cdot & {}^7Li^+ \\ {}^7Li^+ & \cdot\mathcal{E}\cdot & {}^7Li^+ \end{array}\right\} \\ \hline \end{array} \tag{36}$$

- From well established experimental proofs ([Otsu89, Otsu93]) to more recent works ([Warnau2020]), it appears that (i) no sodium-sodium exchange occurs and that (ii) only ”matching” pairs of monomers are active. Accordingly, we assume that these behaviours stand for the lithium cases. Then we discover that $\mathbf{64}$ species are ”blocked” and act only as inhibitors.

- We have $\mathbf{4}$ ”forward” species with a flip rate $f_6$ (respectively $f_7$) when a ${}^6Li^+$ (respectively ${}^7Li^+$) is exchanged:

$$\begin{array}{|c|c|} \hline \text{species} & \text{flip rates} \\ \hline \left\{\begin{array}{ccc} H^+ & \cdot\mathcal{E}\cdot & {}^6Li^+ \\ H^+ & \cdot\mathcal{E}\cdot & {}^6Li^+ \end{array}\right\} & \begin{pmatrix} f_6 \\ f_6 \end{pmatrix} \\ \hline \left\{\begin{array}{ccc} H^+ & \cdot\mathcal{E}\cdot & {}^6Li^+ \\ H^+ & \cdot\mathcal{E}\cdot & {}^7Li^+ \end{array}\right\} & \begin{pmatrix} f_6 \\ f_7 \end{pmatrix} \\ \hline \left\{\begin{array}{ccc} H^+ & \cdot\mathcal{E}\cdot & {}^7Li^+ \\ H^+ & \cdot\mathcal{E}\cdot & {}^6Li^+ \end{array}\right\} & \begin{pmatrix} f_7 \\ f_6 \end{pmatrix} \\ \hline \left\{\begin{array}{ccc} H^+ & \cdot\mathcal{E}\cdot & {}^7Li^+ \\ H^+ & \cdot\mathcal{E}\cdot & {}^7Li^+ \end{array}\right\} & \begin{pmatrix} f_7 \\ f_7 \end{pmatrix} \\ \hline \end{array} \tag{37}$$

- We have $\mathbf{4}$ ”reverse” species with a flip rate $r_6$ (respectively $r_7$) when a ${}^6Li^+$

(respectively $^{7}Li^{+}$) is exchanged:

| species | flip rates |
|---|---|
| $\left\{\begin{array}{ccc} ^{6}Li^{+} & \cdot\mathcal{E}\cdot & H^{+} \\ ^{6}Li^{+} & \cdot\mathcal{E}\cdot & H^{+} \end{array}\right\}$ | $\begin{pmatrix} r_6 \\ r_6 \end{pmatrix}$ |
| $\left\{\begin{array}{ccc} ^{6}Li^{+} & \cdot\mathcal{E}\cdot & H^{+} \\ ^{7}Li^{+} & \cdot\mathcal{E}\cdot & H^{+} \end{array}\right\}$ | $\begin{pmatrix} r_6 \\ r_7 \end{pmatrix}$ |
| $\left\{\begin{array}{ccc} ^{7}Li^{+} & \cdot\mathcal{E}\cdot & H^{+} \\ ^{6}Li^{+} & \cdot\mathcal{E}\cdot & H^{+} \end{array}\right\}$ | $\begin{pmatrix} r_7 \\ r_6 \end{pmatrix}$ |
| $\left\{\begin{array}{ccc} ^{7}Li^{+} & \cdot\mathcal{E}\cdot & H^{+} \\ ^{7}Li^{+} & \cdot\mathcal{E}\cdot & H^{+} \end{array}\right\}$ | $\begin{pmatrix} r_7 \\ r_7 \end{pmatrix}$ |

(38)

#### 2. *Thermodynamic Reduction*

The determination of each $K_{ijkl}$ would require a tremendous amount of work, far beyond the scope of the present manuscript. Indeed, one should vary each concentration, all things equal otherwise, to be able to deduce these values: this is almost impossible because of the very symmetry of the complex.
Nonetheless, we may consider that each site is not a single chemical bond, but rather an electrostatic cavity ready to accept a matching cation.
From the preceding considerations, and for the sake of simplicity and clarity, we shall set:

- for **all** $i,j,k,l$ **of non-futile** species:

$$K_{ijkl} = K_0 \text{ in mol}^{-5} \cdot L^5 \tag{39}$$

- and for all $i,j,k,l$ **futile or blocked** species, $K_{ijkl} \simeq 0$. We hence report the role of unknown $73$ remaining constants on apparent reverse rates, which appears to be sufficient for this model.

#### 3. *Choice and estimation of parameters*

We focused on two curves, measured with 15mM of total lithium. Obviously, we kept this model as simple as possible, but still we have too many unknown parameters. Now, we look for a set of parameters:

- *Qualitative Results*. The adjusted curves are shown in the manuscript, (on Figure 3a for the total intake and Figure 3b for the fractionation) and show that we designed a quantitative and qualitative model. We find a very robust value for $\eta_f$:

$$\eta_f = \frac{V_6^f}{V_7^f} = 1.0148 \pm 0.0004, \ \frac{V_7^f}{V_6^f} = 0.9854 \pm 0.0004 \tag{41}$$

  which means, using Eq. (8), that NHE selects $^6Li^+$ about 4 times more effectively that the electro-osmotic part, and 6.5 times more effectively than the simple diffusion.

- *Quantitative Results*. But we also find out a major interest consisting in the tremendous amount of lithium intake, since a single cell is loaded with about 100mM of total, enriched lithium in a few minutes for only 15mM of external concentration. Compared to a ”passive” NHE-null cell, a simple NHE1 loaded cell loads more than 10 times the amount of $^6Li^+$ in a few minutes, which is remarkable and unprecedentedly reported.

- that allow us to reproduce the overall morphological shapes for both curves,
- and that insists on the robustness of the initial fractionation.

We proceeded as follow.

- We fix the electro-osmotic part computed from the NHE-null cells.
- We use Eq.(33) *sqq* to impose the intracellular proton concentration.
- We fix $K_0 \simeq 10 \cdot 10^{-6}\,\text{mol}^{-5} \cdot L^5$ which is an average value that allows to reproduce the right shapes.
- We start from an apparent maximum rate $V_7^f = f_7 E_0 \simeq 2\,\text{mM.s}^{-1}$.
- We introduce the coefficient $\eta_f$ such that the apparent maximum rate for $^6Li^+$ is:

$$V_6^f = \eta_f V_7^f, \tag{40}$$

  meaning that $\eta_f$ is (i) the forward isotopic speedup of NHE for $^6Li^+$, and (ii) the ”primary” constant we are looking for.
- We symmetrically use $V_7^r$ and $V_6^r = \eta_r V_7^r$ to describe the apparent reverse maximum rate, that on the one hand exists experimentally and helps to expel the huge lithium intake, but that on the other hand is employed to mimic a part of the inhibitor species as described above.
- Since we cannot simultaneously measure the concentration and the fractionation at a given time for the same cell, and since there is a variability between experiments, we scale the computed total lithium concentration with a proportionality factor.
- We adjust the curves with the previously designed algorithm, alternating between the total (quantitative) lithium intake and the (qualitative) corresponding $\delta^7Li^+$ up to the desired convergence.
